# Modeling of Deformable Cell Separation in a Microchannel with Sequenced Pillars

**DOI:** 10.1101/2021.03.01.433451

**Authors:** Scott J. Hymel, Hideki Fujioka, Damir B. Khismatullin

## Abstract

Embedded pillar microstructures are an efficient approach for controlling and sculpting shear flow in a microchannel but have not yet demonstrated to be effective for deformability-based cell separation and sorting. Although simple pillar configurations (lattice, line sequence) worked well for size-based separation of rigid particles, they had a low separation efficiency for circulating cells. The objective of this study was to optimize sequenced microstructures for separation of deformable cells. This was achieved by numerical analysis of pairwise cell migration in a microchannel with multiple pillars, which size, longitudinal spacing, and lateral location as well as the cell elasticity and size varied. This study revealed two basic pillar configurations optimized for deformability-based separation: 1) “duplet” that consists of two closely spaced pillars positioned far below the centerline and above the centerline halfway to the wall; and 2) “triplet” composed of three widely-spaced pillars located below, above and at the centerline, respectively. The duplet configuration is well suited for deformable cell separation in short channels, while the triplet or a combination of duplets and triplets provides even better separation in long channels. These optimized pillar microstructures can dramatically improve microfluidic methods for sorting and isolation of blood and rare circulating tumor cells.

## Introduction

Microfluidic cell separation and isolation is a promising sample-sparring approach for detection of abnormal blood cells or rare circulation tumor cells (CTCs) [1–6]. These diseased cells are often different in their mechanical properties from their circulating counterparts as, e.g., in malaria, sickle cell disease, leukemia, or cancer [1, 2, 7–9], and their isolation requires using deformability-based separation methods. Passive separation of deformable cells holds the most promise for blood diagnostics because it is relatively inexpensive, label-free, and least destructive to the cells than the existing active methods [10–12]. In this approach, cell separation is achieved based on hydrodynamic principles, and it can be controlled by microchannel geometry, inlet flow conditions, or placing embedded structures inside the channel [13–20]. Significant theoretical work is necessary to optimize the passive microfluidic systems for efficient isolation of specific circulating cells.

Embedded microstructures used for size-based cell separation either contained a large number of pillars organized as a symmetric or asymmetric lattice or had a few pillars placed linearly along the flow. The former approach, referred to as deterministic lateral displacement (DLD) [13–15, 21–28], was demonstrated to be much more effective than traditional microfiltration techniques [29–33] because of forcing the cells of specific size to follow a particular streamline instead of trapping large cells. The major issue with the DLD method was clogging the channel at high cell concentration, leading to reduction in separation efficiency and forcing sample dilution to improve efficiency [13, 15]. Defects in pillar size, shape and height appeared during the fabrication process resulted in structural irregularities within the lattice, which further degraded the ability of the DLD systems to separate cells [22]. In addition, to prevent flow disturbances in space between the pillars and channel side walls, side walls were modified or lattice distortion (unequal spacing between pillars) was introduced in the current DLD configurations [24–28].

The above mentioned issues can be overcome by using a sequence of pillars instead of a lattice. Single pillar has been used as a flow divider or for size-based separation of the cells located near the flow centerline [34–36]. With this approach, successful separation of 10, 35 and 120 μm rigid beads were achieved [35]. The numerical simulation revealed that the separation of deformable cells can be achieved by placing a single pillar with a semi-circular cross-section at the flow centerline [36]. Flow sculpting in microchannels with a sequence of pillars placed at different transverse positions have been extensively studied [17, 37–42], but these systems were not shown to be effective for cell-to-cell separation [17, 38, 39]. In particular, complex flow geometries with flow streams moved and split, solution exchange, and separation of embedded identical particles from the solution were created in pillared channels [17]. Sollier *et al.*[39] studied the effect of the spatial arrangement of pillars on separation of white blood cells from lysed blood, but the best they achieved was still less than the separation efficiency for 10 μm rigid particles [17]. Neither experimental nor computational work was done on deformable cell-to-cell separation in a channel with multiple pillars.

Here, we have applied our custom computational fluid dynamics (CFD) algorithm to find sequences of pillars optimal for deformability-based cell sorting. Single and two-cell migration through a sequence of one, two, or three pillars with varying cell deformability, pillar size, and longitudinal and transverse spacing of pillars were investigated. The efficiency of cell-to-cell separation in channels with an optimal three-pillar geometry was tested by a two-cell study in which the cell size, shear elasticity, and initial location were randomized.

## Results and discussion

For size- or deformability-based cell separation in a microchannel, the following criteria need to be satisfied: 1) the transverse separation distance should be large enough to trap the cells with different properties at the channel outlet; and 2) the transverse separation distance should be an increasing function of the cell-to-cell diameter and/or shear elasticity ratios. The former criterion is well established, but a little attention was placed on the latter, which has a key role in cell separation. If the separation distance is large but decreases with the ratio of mechanical properties (“negative trend”), identical cells will be separated, while the cells with different properties will be mixed near the flow centerline.

We first identified these criteria for the single-pillar configuration through the numerical simulation of the pairwise motion of rigid cells of varying diameter and deformable cells of varying elasticity. Figure 1 shows the rigid cell data in the cases of no pillar (NP) and a single pillar which diameter D_p_ ranged from 60 to 150 μm. The pillar was placed near the channel walls (largest absolute value of Y_p_), in the middle between one of the channel walls and the centerline, and at the centerline (Y_p_ = 0 μm). The initial cell-to-cell transverse distance Y_c0_ was fixed at 7 μm (cf. Experimental Section). When the pillar diameter was small (D_p_ = 60 μm), cell focusing, i.e., a reduction in the transverse separation distance occurred for any pillar position (Figure 1a). The focusing effect was strongest for the pillar located at the centerline (Y_p_ = 0 μm). With a slightly larger pillar diameter (90 μm), the separation effect, i.e., an increase in the transverse separation distance, was observed only for a pillar located far above the centerline (Y_p_ = 55 μm, Figure 1b). At this position, the motion of the first cell but not the second one was influenced by the pillar. If the pillar was large (120 or 150 μm), cells split by the centrally located pillar, leading to a large separation distance between the cells (Figure 1c,d). The separation distance increased by 28 μm after passing the 150 μm pillar located at Y_p_ = 0 μm (Figure 1d). Other configurations caused the focusing effect.

**Figure 1:**
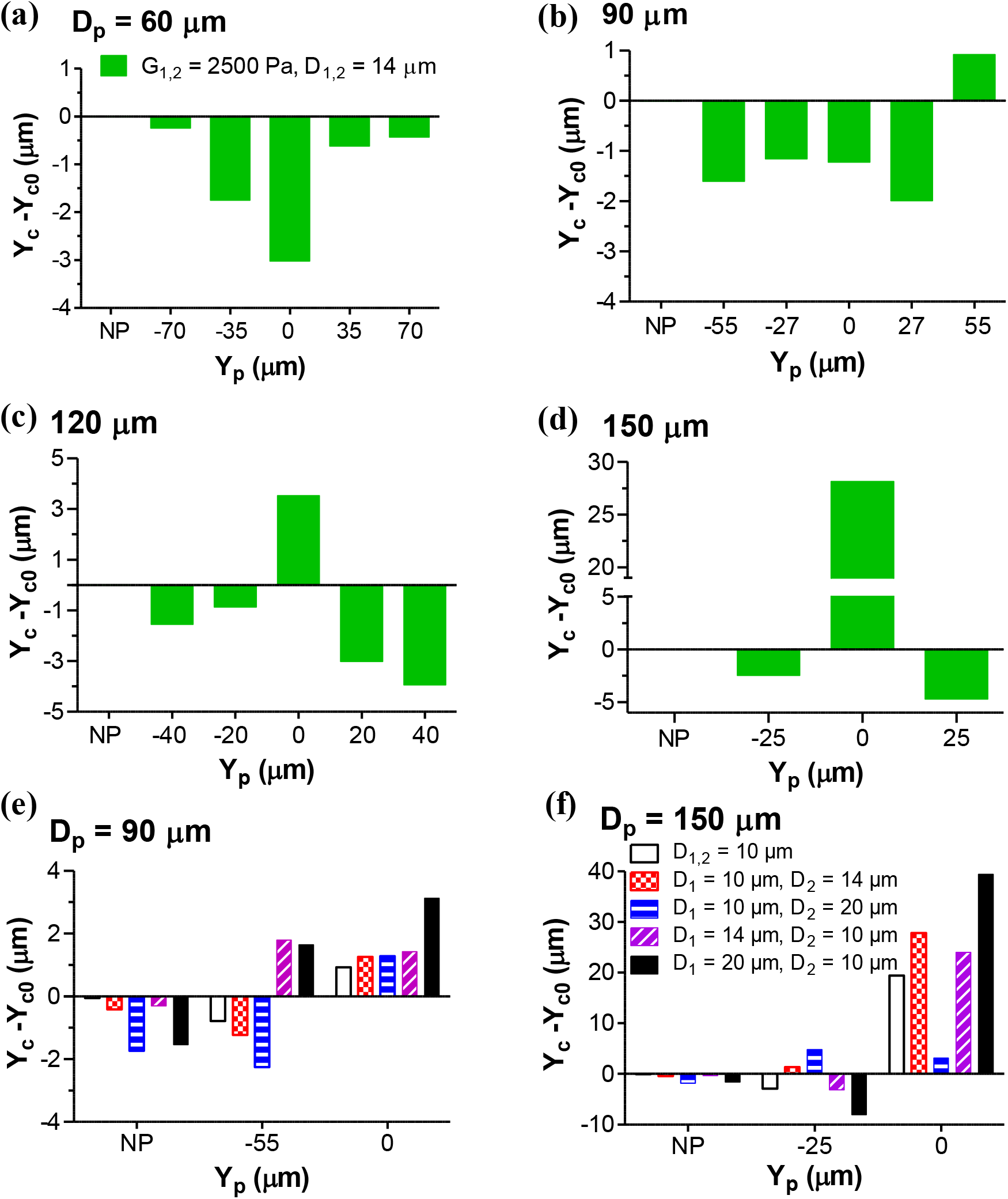
Change in transverse separation distance between two rigid cells migrated through a microchannel with a single pillar, as a function of pillar location Y_p_. (a-d) Effect of pillar diameter D_p_ on separation of identical rigid cells. (e,f) Effect of pillar diameter on separation of different size rigid cells. The shear elasticity of rigid cells is 2500 Pa. NP represents a microchannel without a pillar.

The cells of different size were studied for pillar diameters of 90 and 150 μm. Two scenarios were considered: 1) cell #1, located slightly above the centerline, was smaller than cell #2, located below the centerline (in this case, D_2_/D_1_ was the cell diameter ratio); and 2) cell #1 was larger than cell #2, with D_1_/D_2_ being the cell diameter ratio. Without a pillar, the focusing effect, more pronounced at a larger cell diameter ratio, was observed for both scenarios (Figure 1e,f). If a 90-μm pillar was placed along the bottom wall (Y_p_ = −55 μm, Figure 1e), cell focusing observed for D_1_ < D_2_ switched to cell separation with a negative trend with respect to the cell diameter ratio for D_1_ > D_2_. With a large, 150 μm pillar near the bottom wall (Y_p_ = −25 μm, Figure 1f), the cell behavior changed again between the two scenarios. If D_1_ < D_2_, cells were separated, and the separation distance slightly increased with the cell diameter ratio. If D_1_ > D_2_, the cells were focused. For a pillar positioned at the centerline, the separation effect occurred for all cell diameters. However, the second criteria for cell separation (higher effect for larger cell diameter ratio) was not always satisfied. If D_p_ = 90 μm (D_p_/W = 0.45), both the criteria were met for D_1_ > D_2_ or D_1_ < D_2_ (Figure 1e). Despite a large separation distance achieved for D_p_ = 150 μm (D_p_/W = 0.75) in the first scenario, there was a drop in the separation distance by 36.3 μm between D_2_/D_1_ = 1.4 and 2.0 (Figure 1f). No such drop was observed in the second scenario. Thus, with central placement of the pillar, size-based separation requires that larger cells are located above the centerline while smaller cells are below the centerline before entering the pillar space. Previous experiments with pillars of half the channel width (D_p_/W = 0.5) demonstrated 50% and 95% trapping efficiency for deformable cells (leukocytes) and 10 μm rigid particles, respectively [17, 39]. None of the experimental work, however, demonstrated separation of cells or particles of multiple types/sizes using a single pillar or a linear sequence of pillars. It is unclear if the central positioning of the pillar will be effective for separation of cells with different deformability.

Figure 2 shows the effect of cell-to-cell elasticity ratio on the cell separation in a channel without a pillar or with a single pillar. To reduce the simulation time, the central location of the pillar was at Y_p_ = −3 μm instead of 0 μm. In the absence of the pillar, the transverse separation distance increased by less than 0.30 μm and was slightly dependent on the elasticity ratio (Figure 2). The separation distance between the cells grew by 0.37 to 0.85 μm when they passed a 60 μm size pillar (D_p_/W = 0.3) located far from the centerline (Y_p_ = ±70 μm) (Figure 2a,e). When the pillar was close to the centerline, cell focusing was observed. The increase in the pillar diameter amplified the cell focusing effect (Figure 2b-e). For example, the transverse separation distance reduced to −0.9 μm (cells swapped) post a 150 μm pillar (D_p_/W = 0.75) for the cells with elasticity of 50 and 500 Pa (Figure 2d). The rigid-cell separation was only observed for a 90-μm pillar located far above the centerline (Figure 2b). To summarize, a slight separation of deformable cells can be achieved in a microchannel with a single small-diameter pillar positioned away from the centerline.

**Figure 2:**
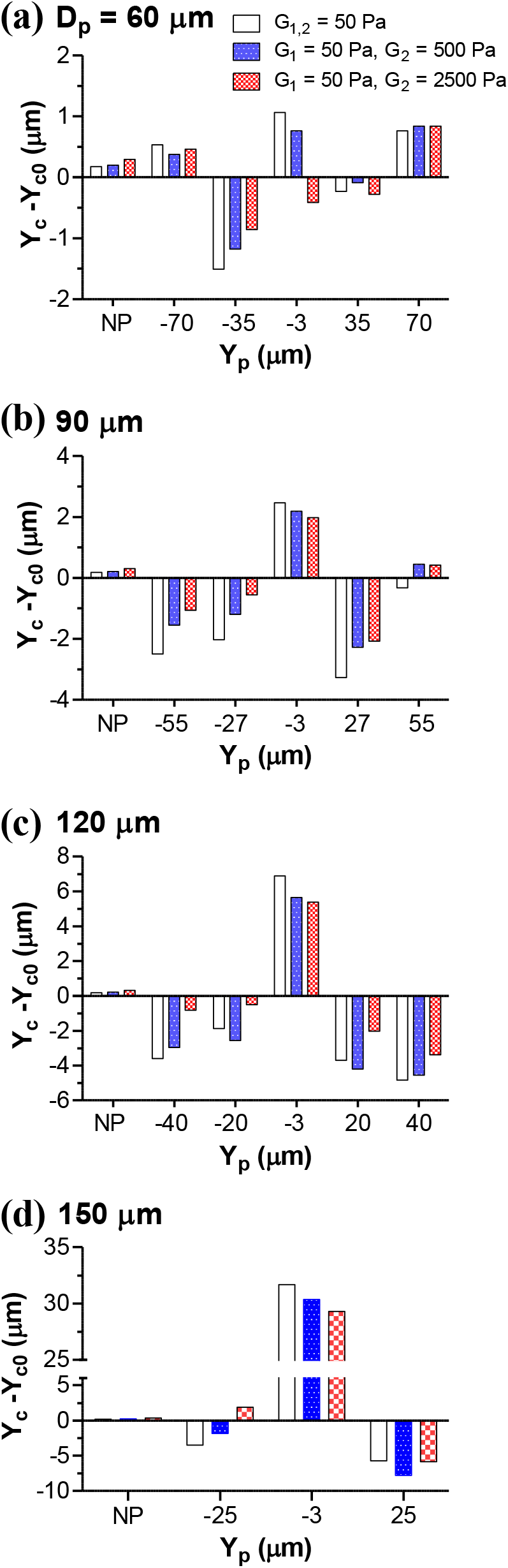
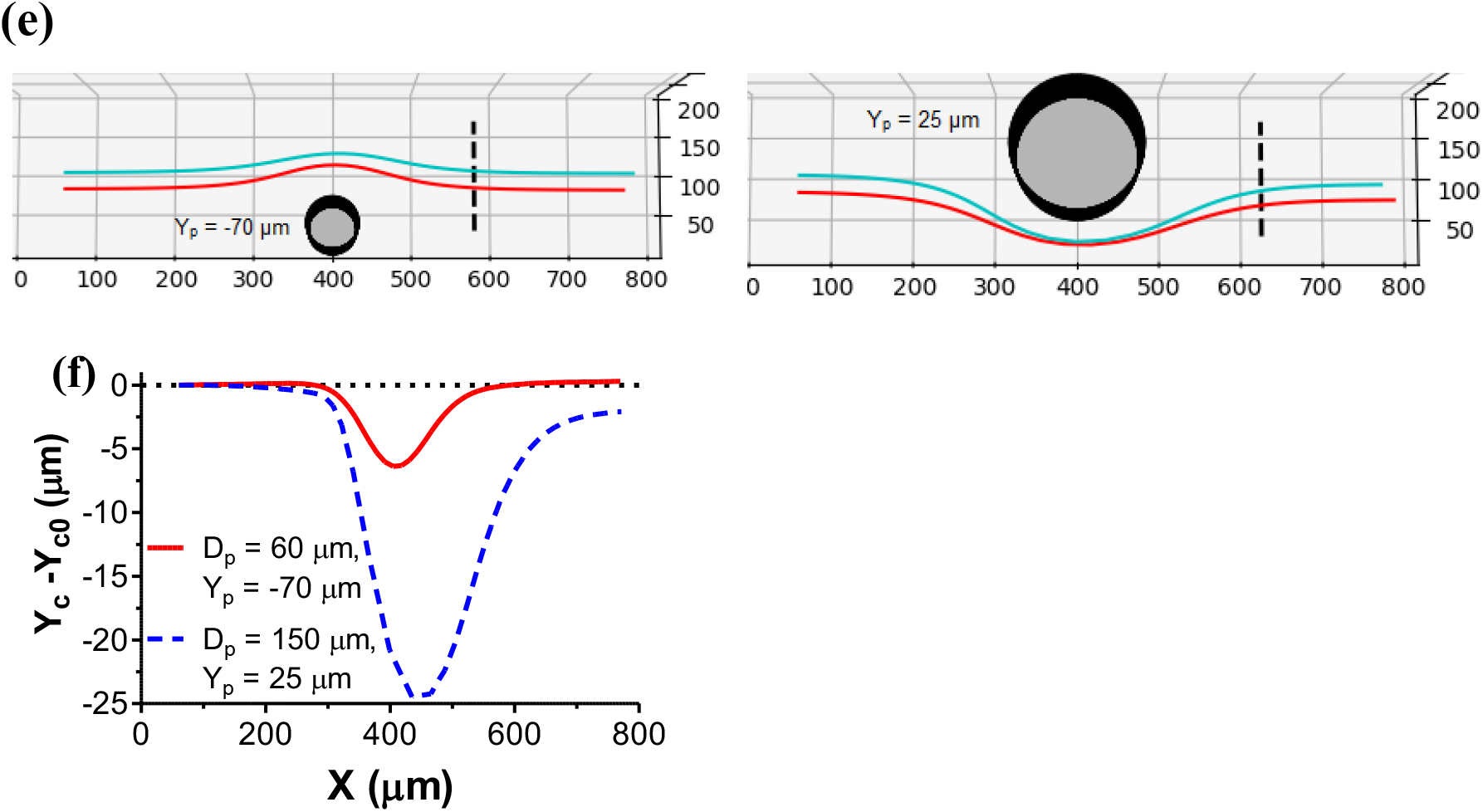
Migration of two 14-μm diameter cells with equal or unequal shear elasticity through a single pillar microchannel. (a-d) Change in transverse separation distance between the cells as a function of pillar location for different pillar diameters. NP represents a microchannel without a pillar. (e) Trajectories of the cells in a microchannel with a thin pillar (D_p_ = 60 μm) located far below the centerline (Y_p_ = −70 μm, left) and with a thick pillar (D_p_ = 150 μm) above the centerline (Y_p_ = 25 μm, right). Here, cells #1 (cyan) and #2 (red) have a shear elasticity of 50 Pa and 2500 Pa, respectively. (f) A thinner pillar, depending on its location with respect to the cells, offers a more positive change in the transverse separation distance between the cells with unequal elasticity than a thicker pillar does.

We next studied the separation of deformable cells in a channel containing two 60-μm diameter pillars at different locations and inter-pillar spacing (S_p_) from 30 to 180 μm (Figure 3). There was a distinct behavior for small and large spacing. First, when S_p_ = 30 or 60 μm, both the criteria for deformability-based cell separation were satisfied when the first and second pillars were located far below and above the centerline, respectively (Y_p1,2_ = −70, 35 μm, Figure 3a,b). With S_p_ = 30 μm and the first cell being highly deformable (G_1_ = 50 Pa), Y_c_ – Y_c0_ increased from 1.5 to 3.3 μm with an increase in elasticity of the second cell G_2_ from 50 to 2,500 Pa (white vs red bars in Figure 3a; see also trajectories in Figure 3f). The separation effect diminished and then disappeared at higher S_p_ values for this configuration of pillars (Figure 3b-d). Moving the first pillar within close proximity of the centerline (Y_p1_ = −3 μm) led to either a strong focusing effect or a negligible separation effect for different inter-pillar spacings. Moving the second pillar to far above the centerline (Y_p2_ = 70 μm) resulted in the negative trend with respect to the cell elasticity ratio. These data indicate that far below and at 50% above the centerline could be optimal locations of first and second pillars, respectively, for the closely spaced “duplet” configuration.

**Figure 3:**
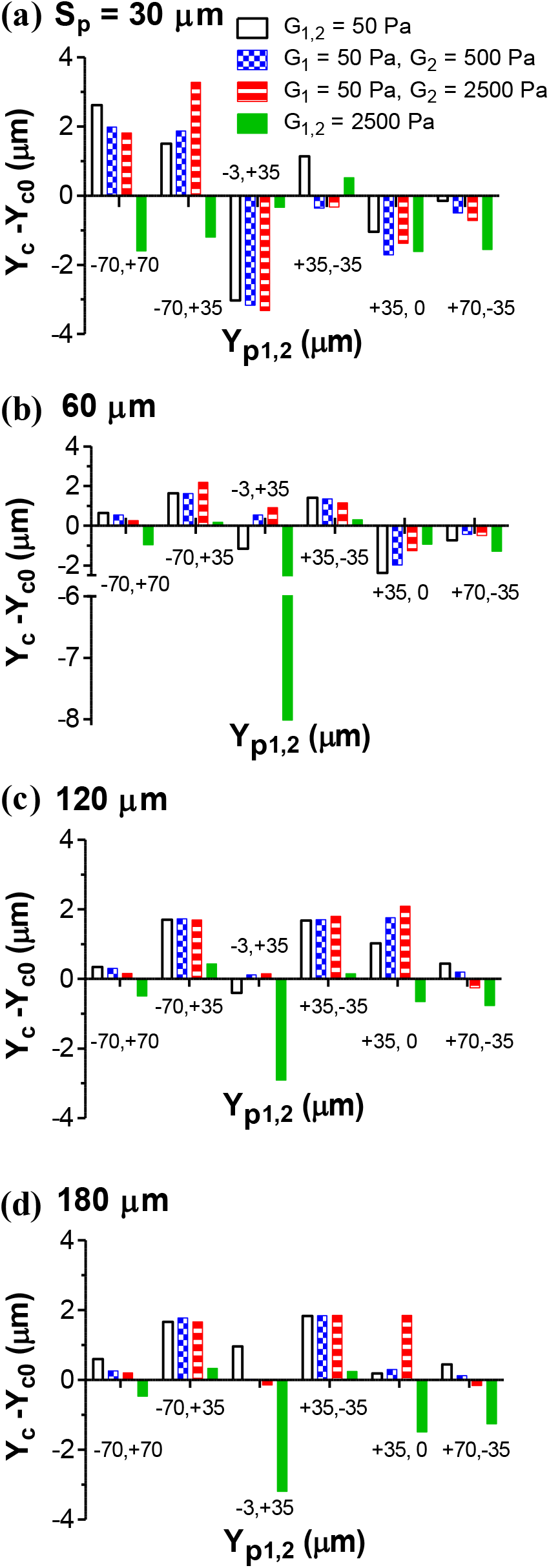
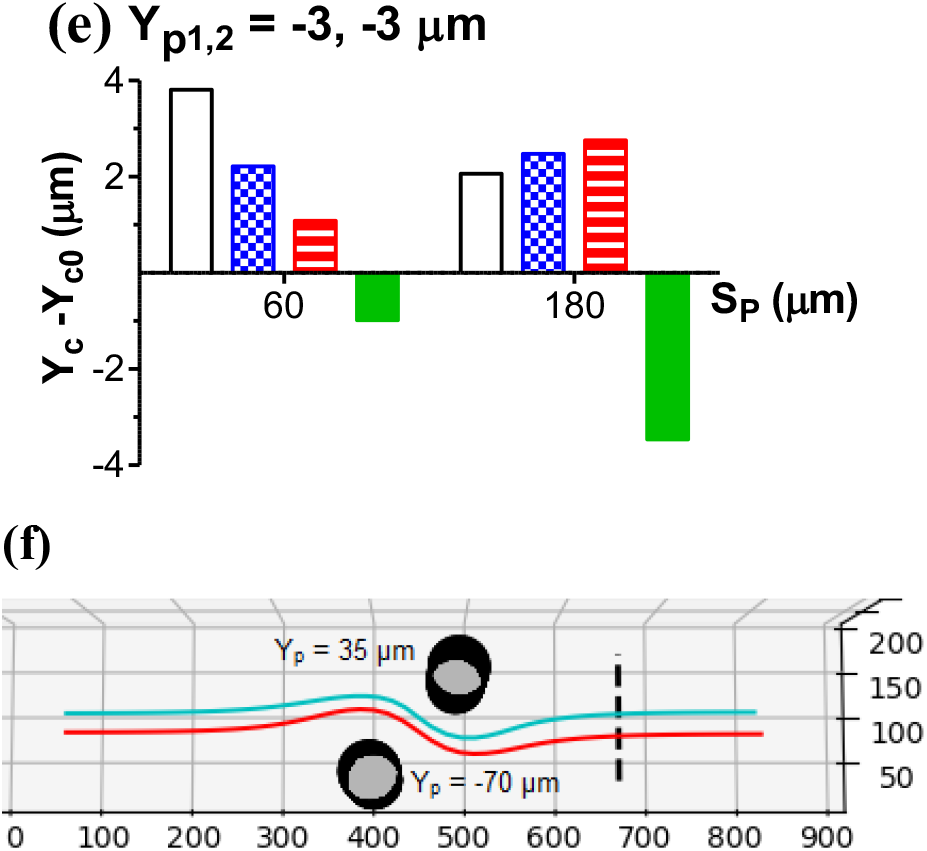
Change in transverse separation distance between two 14-μm diameter cells with equal or unequal elasticity migrated through a microchannel with two pillars, as a function of pillar location. (a-d) Effect of longitudinal spacing, Sp, of 60-μm diameter pillars on cell separation. (e) Transverse separation distance vs. inter-pillar spacing for a line sequence of pillars near the centerline (Y_p1,2_ = −3, −3 μm). (f) Trajectories of the cells in a microchannel with two thin, closely spaced pillars (D_p_ = 60 μm, S_p_ = 30 μm) located far below (Y_p_ = −70 μm) and above the centerline (Y_p_ = 35 μm), respectively. Cells #1 (cyan) and #2 (red) have a shear elasticity of 50 Pa and 2500 Pa, respectively.

Second, for large inter-pillar spacing (120 and 180 um), the separation effect with the positive cell elasticity ratio trend was observed for Y_p1,2_ = 35, 0 μm (Figure 3c,d). This result further indicates that the first pillar should not be located at the centerline. However, placing the second pillar at the centerline may work when the distance from the first pillar is sufficiently large. The configuration at which the sequence of pillars is located in the vicinity of the centerline (Y_p1,2_ = −3, −3 μm), previously tested in experimental work [17, 39, 42], is not viable for efficient deformability-based separation no matter how far or near pillars are spaced (Figure 3e). Note that most of the configurations shown in Figure 3 led to focusing of rigid cells (green bars), while separation was observed for deformable cells in some cases. Thus, the optimization of pillar placement for deformability-based separation should not be based on rigid cell behavior.

The positions of the first two pillars in the three-pillar simulation were selected based on the data obtained in Figs. 2 and 3, specifically, the pillars were selected to be close to the channel walls (±70 μm) or in the middle between the centerline and one of the walls (±35 μm). The third pillar was positioned at Y_p3_ = ±35 μm or 0 μm. All the pillars in this simulation had the same diameter D_p_ = 60 μm. As seen in Figure 4a, when inter-pillar spacing was small (S_p_ = D_p_/2 = 30 μm), the deformability-based cell separation was achieved only for Y_p1,2,3_ = −70,+35,-35 μm. However, the separation effect was not strong: maximum increase over the initial separation distance was only 1.4 μm. For S_p_ = D_p_ (Figure 4b), the separation distance increased, but the negative elasticity effect occurred in most cases. The Y_p1,2,3_ = 35,-35,0 μm configuration led to large deformability-based separation for S_p_ = 2D_p_ (Figure 4c). For the longest spacing studied (S_p_ = 3D_p_, Figure 4d,e), Y_p1,2,3_ = +35,-35, 0 μm and especially Y_p1,2,3_ = −70,+35, 0 μm led to large separation with a strong and positive elasticity effect. Overall, for the widely spaced set of three pillars (“triplet”), positioning of the first pillar far below the centerline, the second pillar above the centerline at half distance to the wall, and the third pillar at the centerline was most efficient for cell separation.

**Figure 4:**
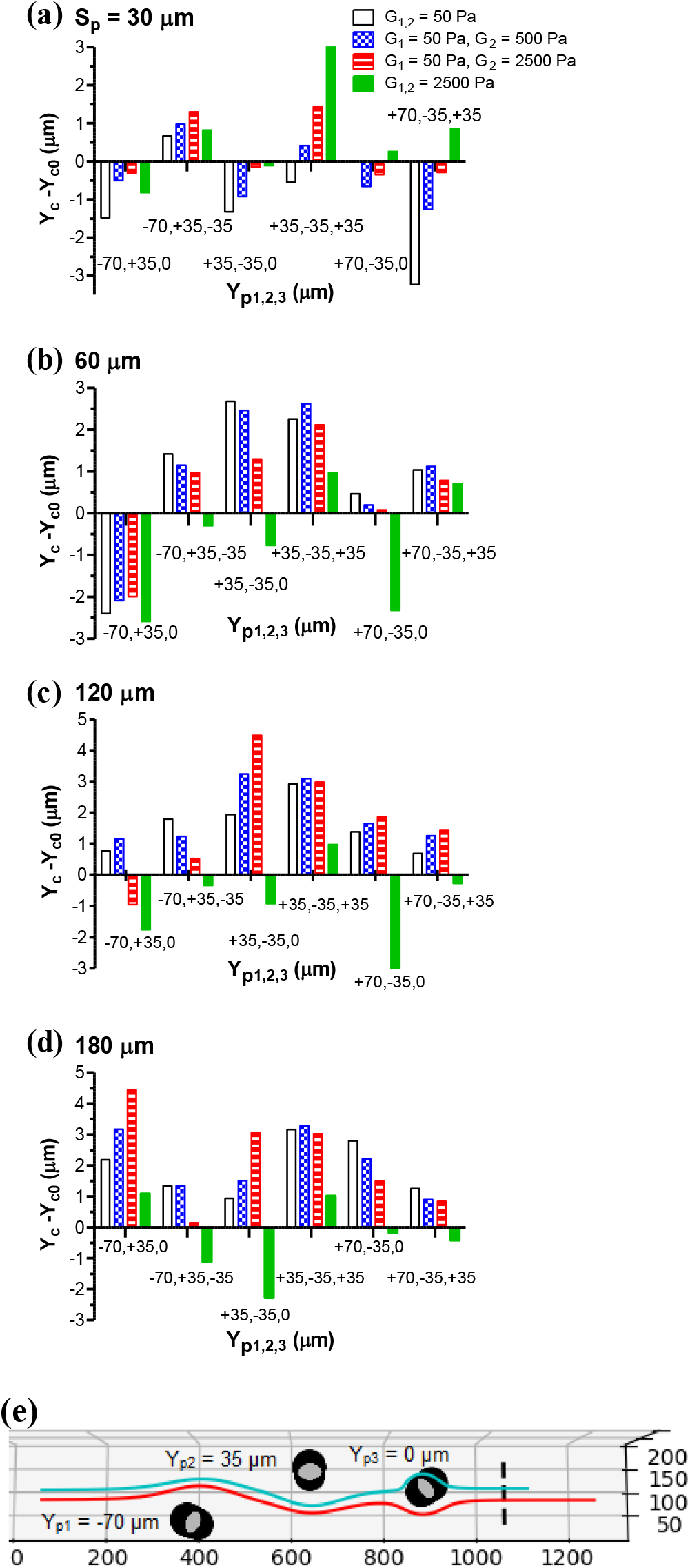
Change in transverse separation distance between two 14-μm diameter cells with equal or unequal elasticity migrated through a microchannel with three pillars, as a function of pillar location. (a-d) (a-d) Effect of inter-pillar spacing on cell separation. (e) Trajectories of the cells for widely spaced pillars (S_p_ = 180 μm) located at Y_p1,2,3_ = −70, +35, 0 μm. The shear elasticity of cell #1 (cyan) and #2 (red) was 50 Pa and 2500 Pa, respectively. The pillar diameter was 60 μm.

Figure 5 demonstrates how the increase in pillar number influenced the cell-to-cell separation distance for closely and widely spaced pillars. A single pillar placed far below the centerline (Y_p1_ = −70 μm) caused a negligible change in the separation distance, independently on the cell elasticity ratio and pillar spacing. Placing the second pillar above the centerline (Y_p2_ = 35 μm) led to strong deformability-based cell separation for small inter-pillar distance (“duplet", Figure 5a,c). Nonetheless, it did not induce a change in the separation distance for cells with different deformability, when the inter-pillar distance was large (Figure 5b,c). Adding the third pillar at the centerline (Y_p3_ = 0 μm) caused a focusing effect in the case of closely spaced pillars (Figure 5a). However, the strong separation effect with a positive deformability ratio trend occurred for widely spaced pillars (Figure 5b,c). For this “triplet” configuration, the separation distance increased even for rigid cells.

**Figure 5:**
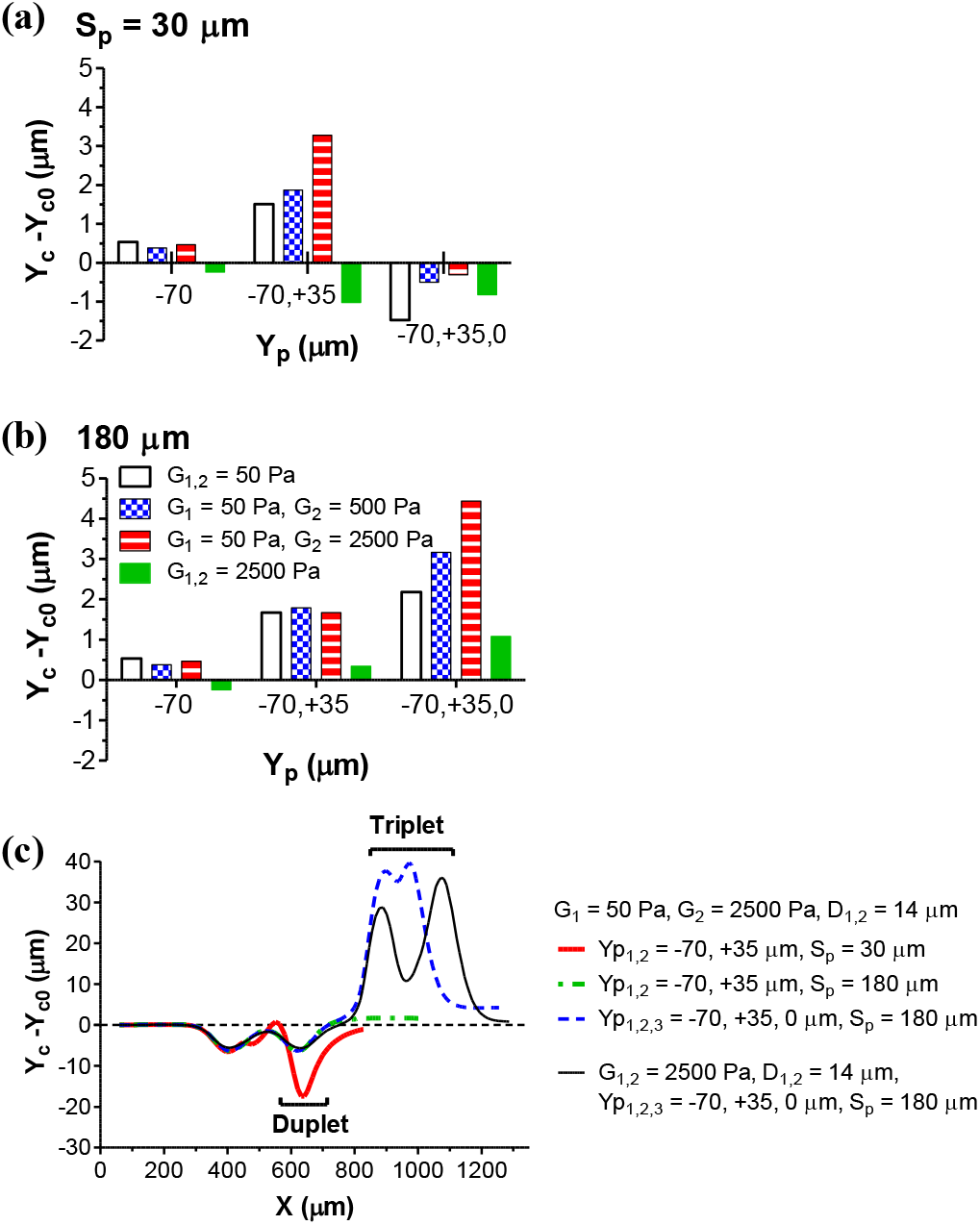
Effect of progressively adding pillars on the transverse separation distance between cells migrated through closely spaced pillars (S_p_ = 30 μm, a) or widely spaced pillars (S_p_ = 180 μm, b). (c) Change in the transverse separation distance between soft (G_1_ = 50 Pa) and rigid cells (G_2_ = 2500 Pa) with the longitudinal migration distance X for “duplet” and “triplet” configurations of pillars (red solid line, blue dashed line). Also shown in (c) is the transverse separation distance of soft and rigid cells for the duplet configuration with large inter-pillar spacing (green dashed line) and the transverse separation distance of rigid cells for the triplet pillar configuration (black line). The pillar and cell diameters were fixed at 60 μm and 14 μm, respectively.

An increase in pillar size is detrimental for deformability-based cell separation when considering off-center configurations of pillars, as evident from both the single-pillar (Figure 2) and three-pillar data (Figure 6). While the Y_p1,2,3_ = 35,-35, 0 μm configuration caused strong separation at D_p_ = 60 μm and S_p_ = 2D_p_ (Figure 6a,b), an increase in the pillar dimeter above 90 μm by keeping the pillar spacing-to-diameter ratio fixed led to an insignificant change in or a negative trend of the separation distance with respect to the cell elasticity ratio (Figure 6a,c).

**Figure 6:**
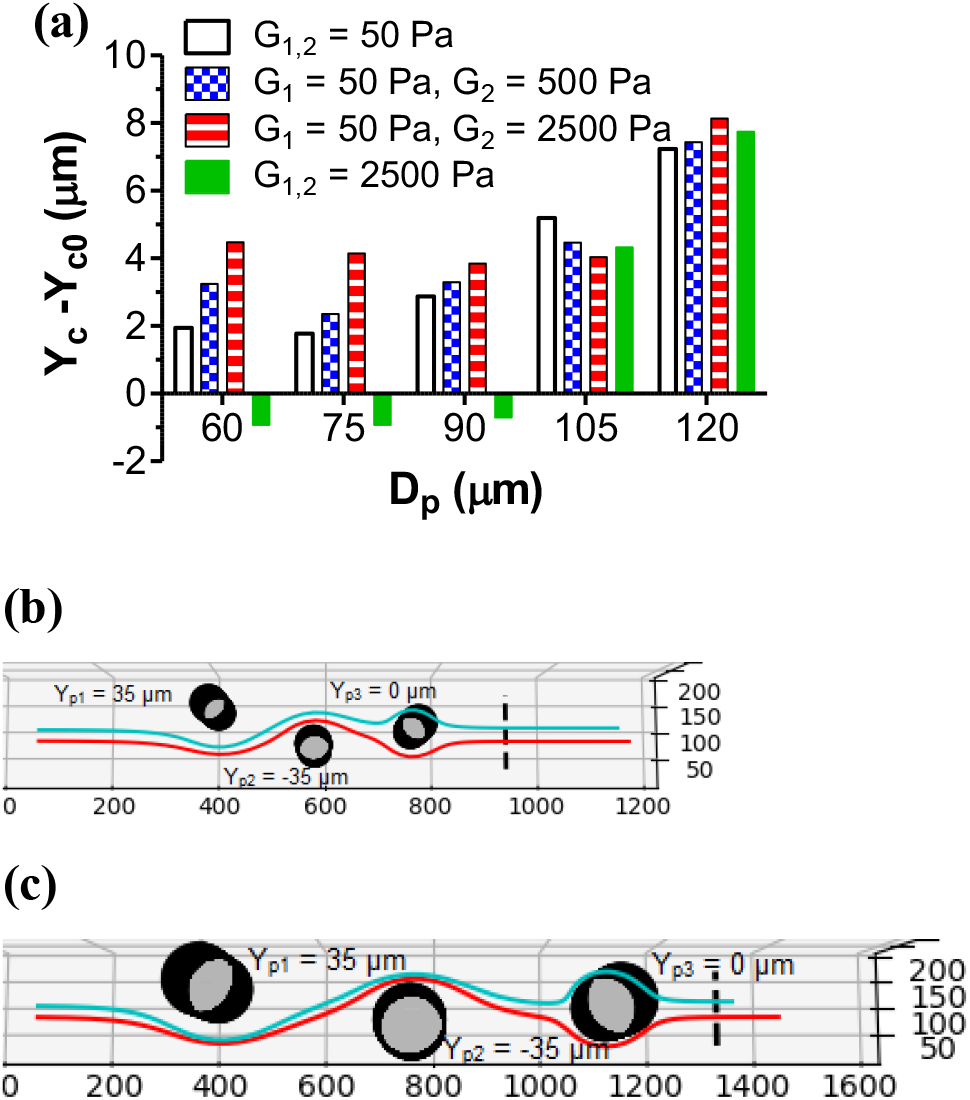
Effect of pillar diameter on the separation of 14-μm diameter cells with equal or unequal elasticity in a microchannel with three pillars. (a) Change in the transverse separation distance vs. pillar diameter for pillars located at Y_p1_ = +35 μm, Y_p2_ = −35 μm, Y_p3_ = 0 μm. Here, the inter-pillar spacing was fixed in relation to pillar diameter (S_p_ = 2D_p_). Also shown are trajectories of soft (G_1_ = 50 Pa, cyan) and rigid cells (G_2_ = 2500 Pa, red) for thin pillars (D_p_ = 60 μm, b) and thick pillars (D_p_ = 120 μm. c).

When the cells were in close proximity to the centerline, two configurations of three pillars led to deformability-based cell separation: Y_p1,2,3_ = 35,-35, 0 μm (Case 1, Figure 7a) and −70,35, 0 μm (Figure 4). We conducted the randomized numerical study to test which of them works better for separation of circulating cells, which location was not restricted to the vicinity of the centerline. In this study, the second configuration was slightly altered to −60,40, −10 μm (Case 2, Figure 7b) to accommodate large variations in cell size and location. The pillar diameter and the inter-pillar spacing was fixed at 60 μm and 120 μm, respectively, for both cases.

**Figure 7:**
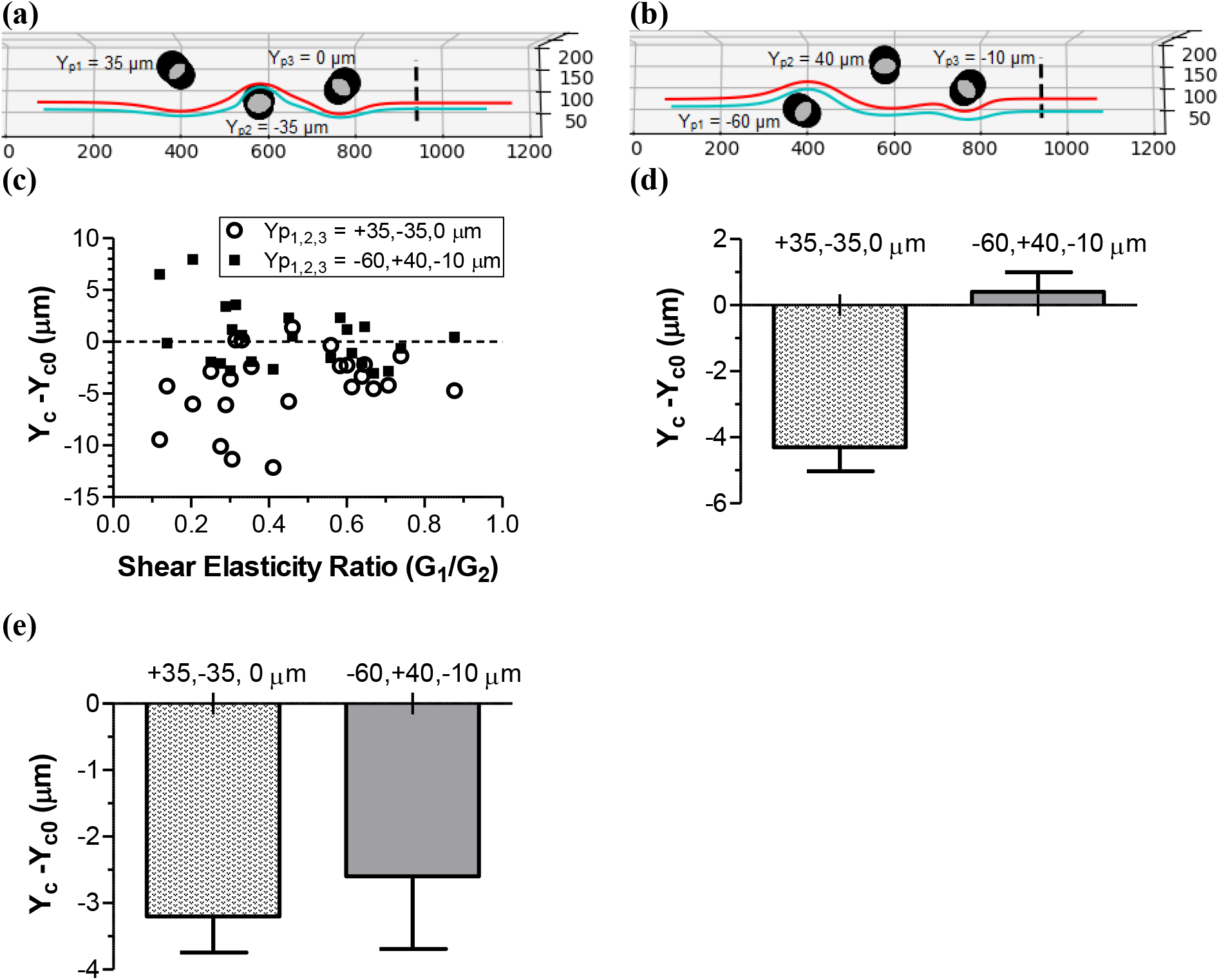
Data of the randomized study for two three-pillar configurations: Y_p1_ = +35 μm, Y_p2_ = −35 μm, Y_p3_ = 0 μm (a, open circles in c, light bar in d and e) and Y_p1_ = −60 μm, Y_p2_ = +40 μm, Y_p3_ = −10 μm (b, filled squares in c, dark bar in d and e). The inter-pillar spacing and pillar diameter were fixed at 120 μm and 60 μm, respectively. The initial location of and distance between the cells as well as the diameter and shear elasticity of the cells were randomly selected. (a, b) Representative trajectories of the cells for above-mentioned configurations of pillars. A more deformable cell (red) was initially located closer to the centerline than a less deformable one (cyan). (c) Change in transverse separation distance between the cells with similar cell diameter but different elasticity values for each of the configuration. (d) Mean ± SEM (N = 24) of the data in (c). (e) Mean ± SEM (N = 8) for the change in transverse separation distance between the cells with different diameter and similar shear elasticity (elasticity ratio of 0.8 or higher) for each of the pillar configurations.

Figure 7(c,d) shows the post-pillar separation distances for 24 out of 50 randomized runs in which the size difference between the cells was 2 μm or less. The data indicate the capability to separate the cells similar in size but differ in deformability. As seen in Figure 7(c), focusing predominantly occurred in Case 1, while more than half the runs led to the separation effect in Case 2. On average, the separation distance was highly negative in Case 1 and positive in Case 2 (Figure 7d). Thus, the 3-pillar configuration at which the first pillar is far below the centerline, the second pillar above the centerline, and the third pillar near the centerline (“triplet”) is most efficient for deformability-based cell separation. Note that both configurations studied led to the focusing effect when the cells have different diameters but the shear elasticity ratio between them were 0.8 or higher (Figure 7e). This indicates the positioning the first two pillars off the centerline is not beneficial for size-based cell separation.

## Conclusion

The results of our study point out to significant differences in pillared microchannel design criteria between size- and deformability-based cell separation. Placing pillars at the flow centerline allows for efficient separation of different size cells with similar mechanical properties, as already tested experimentally [17, 35, 39], while the off-center pillar positioning is better suited for separation of cells with different mechanical properties. In particular, our computational analysis revealed the “duplet” and “triplet” pillar configurations, optimal for separating deformable cells, in which either both pillars or the first two pillars are located away from the flow centerline. In the duplet configuration, longitudinal spacing between pillars is small (half the pillar diameter), i.e., a sequence of duplets can be used to separate cells over short distances. A sequence of triplets or a combination of duplets and triplets leads to a large separation than the duplets only, but it requires the use of a long channel. Based on data in Figs. 1 and 7, mixing the duplet and triplet configurations of pillars with a sequence of centrally located pillars may cause separation of the cells different in both size and deformability.

## Experimental Section

### Simulation Models and Methods

The numerical simulation of deformable cell migration in a microchannel with pillars has been conducted using a custom computational fluid dynamics algorithm referred to as VECAM [43, 44]. This algorithm solves a full set of Navier-Stokes equations for a multiphase fluid using the volume-of-fluid (VOF) method, with each phase being treated as either Newtonian or a viscoelastic (Oldroyd-B) fluid. In the current study, the single-phase model of the cell [7, 45–51] in which the interior of the cell (cytoplasm) was viscoelastic while the extracellular fluid was Newtonian was employed. The cytoplasmic viscoelasticity was described by the following form of the Oldroyd-B model:

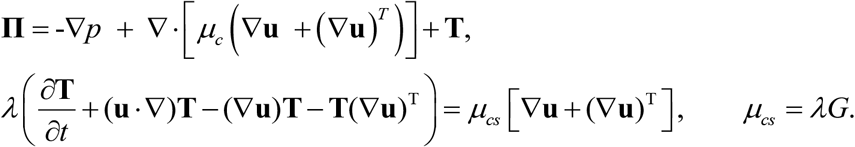

Here **Π** is the total stress tensor that includes the contributions from pressure *p*, cytosolic shear viscosity *μ*_*c*_, and cytoskeletal viscoelasticity determined by tensor **T**; **u** the velocity vector, *ρ* mass density, *μ*_*cs*_ cytoskeletal shear viscosity, *G* cytoskeletal shear elastic modulus, and *λ* cytoplasmic relaxation time.

The geometry of the computational domain was built from the extrusion of bitmap images of channels with one, two, or three pillars (Figure 8a). The channel height and width were fixed at 50 μm and 200 μm, respectively, according to microfluidic systems designed and tested in Di Carlo’s laboratory [17, 39, 42]. The total length varied from 800 μm to 1600 μm depending on the number and diameter of pillars and inter-pillar spacing. Pillars with a circular cross-section were studied, each characterized by the diameter *D*_*p*_ and longitudinal (X) and transverse (Y) positions. Prior to placing the cells in the microchannel, fully developed flow was established (Figure 8b).

**Figure 8:**
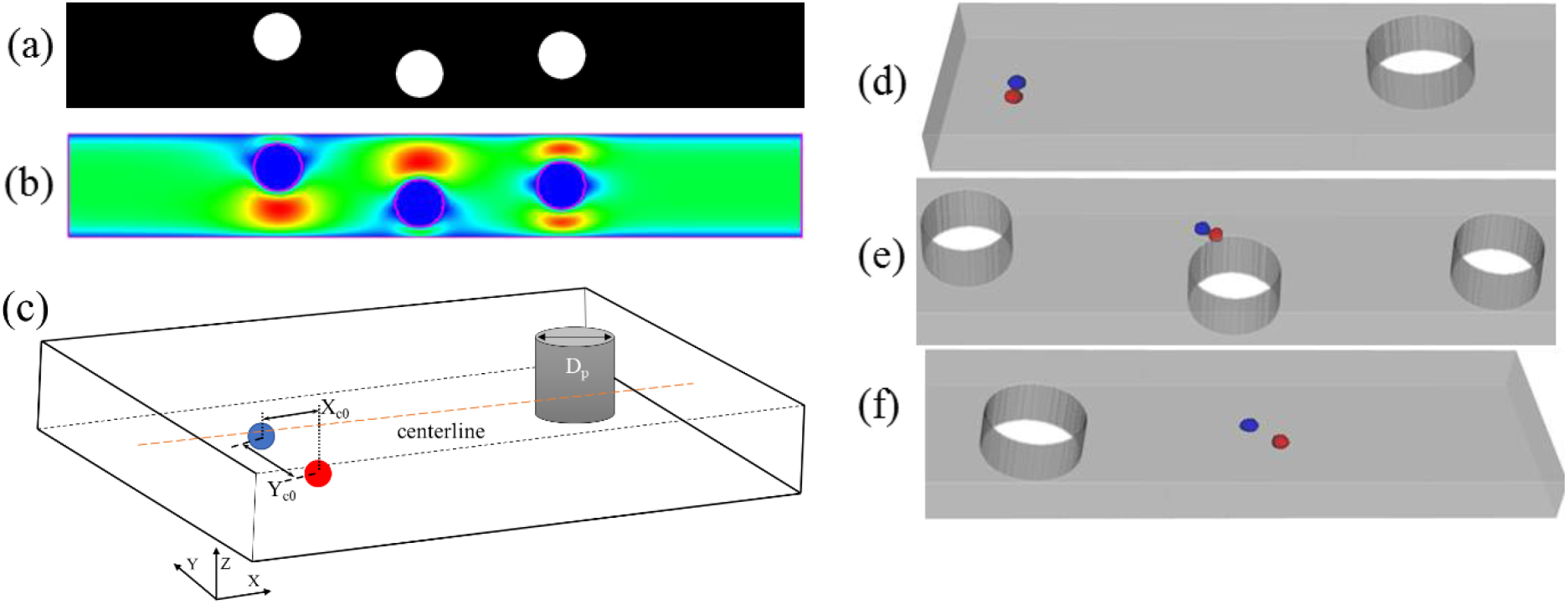
(a) Python bitmap image created for a three-pillar system with each having a pillar diameter of 90 μm. (b) Fully developed flow field for this pillar configuration (slice taken at midplane in Z-direction). (c) Schematic of cell placement demonstrating X_c0_ and Y_c0_ measurements with respect to the channel centerline and in relation to the first pillar and the inlet. (d-f) Soft (G_1_ = 50 Pa, blue) and rigid cells (G_2_ = 2500 Pa, red) moving through a sequence of three pillars (the flow field for this system is shown in b). The change in transverse separation distance was calculated as the difference between the separation distance at 150 μm past the last pillar (f) and the initial separation distance (d).

All the data were produced on a computational mesh with cubic grid elements of 1.0 μm in size (regular mesh). To test if this mesh size provides sufficient accuracy, the simulation of two cells (G_1_ = 50 Pa, G_2_ = 2500 Pa) in motion initially located near the centerline in a single pillar (D_p_ = 60 μm, Y_p_ = 65 μm) was additionally done on the coarse mesh (1.5 μm in size) and the fine mesh (0.5 μm). In this mesh refinement study, there was less than 25% error in the change in transverse separation distance between cells in question when comparing the output of fine and regular mesh. While comparing the output of regular and coarse mesh, there was 128% error in the change in separation between cells after the pillars. Finally, there was an 185% error between the fine and coarse mesh. Based on this data, a 1.0 μm grid has been selected for all the cases.

### Simulation Protocols

In the deterministic study, two cells were placed slightly above the centerline (cell #1) and farther below the centerline (cell #2). The cells were either identical or had different size or deformability. In most runs, cell #1 was more deformable to replicate the conditions for blood flow in microvasculature and microfluidic systems (deformable RBCs near the centerline) [52–55]. The cells were initially located at least 250 μm from the outer edge of the first pillar and at least 60 μm from the inlet. They were separated transversely by distance Y_c0_ = 7 μm, as shown in Figure 8c. The initial cell diameter *D* and shear elasticity *G* varied from 10 to 20 μm and 50 to 2,500 Pa, respectively. The relaxation time of the cell was fixed at 0.17 s [1, 7]. The pillar diameter (D*p*) ranged from 60 to 150 μm. The transverse positions of pillars (Y_p_) ranged from –70 μm to +70 μm, with central positioning at Y_p_ = 0 μm. Changes in between the outer edges of the cells were measured and recorded as the cells migrate through each sequence of micropillars (Figure 8d-f) at different time instances to assess the effect of pillars on cell separation. The differences between the transverse separation distance at 150 μm after the last pillar and the initial separation distance have been reported (Figure 1e). The cells reached steady state at a longitudinal distance of 150 μm from the last pillar.

In the randomization study, the cells were initially located at 60 to 100 μm away from the inlet in the X-direction, between −50 and 50 μm from the centerline in the Y-direction, and at 25 μm (mid-plane) in the Z-direction. These restrictions were used to accommodate different cell sizes. The total separation distance, 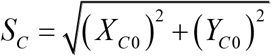 (Figure 8c), varied between 3 and 12 μm. while cytoskeletal shear elasticity and cell diameter ranges were 100 to 3500 Pa and 10 to 24 μm, respectively. The uniform distribution was applied to randomize the location of and separation distance between the cells, while the normal distribution was used for the cell diameter and shear elasticity. In the latter, the mean ± standard deviation was 17 ± 4 μm for the cell diameter and 1200 ± 500 Pa for the shear elasticity. We selected the properties of these groups by considering 24 different types of cells including healthy epithelial cells and carcinoma cells from several tissues (breast, lung, prostate, ovaries, bladder) as well as healthy and malignant leukocytes (leukemia, lymphoma) [4, 56–68]. A total of 50 randomized simulations were conducted for each of the pillar configurations.

## Acknowledgements

The authors acknowledge funding from National Science Foundation (grant No. 1301286) and Louisiana Board of Regents (LEQSF(2011-14)-RD-A-24). This work was supported in part using high performance computing (HPC) resources and services provided by Technology Services at Tulane University, New Orleans, Louisiana. It also used the Bridges system, which is supported by National Science Foundation grant number ACI-1445606, at the Pittsburgh Supercomputing Center (PSC). The authors thank Dino Di Carlo for helpful discussion.

## Conflicts of Interest

There are no conflicts to declare.

## Notes

### Competing Interest Statement

The authors have declared no competing interest.

